# The Potential of Low Press and Hypoxia Environment in Assisting Pan-cancer Treatment

**DOI:** 10.1101/2023.03.23.534056

**Authors:** Xiaoxi Hu, Xinrui Chen, Mengzhen Sun, Xilu Wang, Zixin Hu, Shixuan Zhang

**Author notes:** Corresponding author: Xilu Wang’s; Zixin Hu’s; Shixuan Zhang’s. These authors contributed equally to this work (Xiaoxi Hu, Xinrui Chen, Mengzhen Sun).

## Abstract

**Objective:** A low incidence and mortality rate of cancer has been observed in high-altitude regions, suggesting a potential positive effect of low press and hypoxia (LPH) environment on cancer. Based on this finding, our study aimed to construct a pan-cancer prognosis risk model using a series of ADME genes intervened by low oxygen, to explore the impact of LPH environment on the overall survival (OS) of various kinds of cancers, and to provide new ideas and approaches for cancer prevention and treatment.

**Datasets and Measures:** The study used multiple sources of data to construct the pan-cancer prognosis risk model, including gene expression and survival data of 8,628 samples from the cancer genome atlas, and three gene expression omnibus databases were employed to validate the prediction efficiency of the prognostic model. The AltitudeOmics dataset was specifically used to validate the significant changes in model gene expression in LPH. To further identify the biomarkers and refine the model, various analytical approaches were employed such as single-gene prognostic analysis, weighted gene co-expression network analysis, and stepwise cox regression. And LINCS L1000, AutoDockTools, and STITCH were utilized to explore effective interacting drugs for model genes.

**Main Outcomes and Conclusions:** The study identified eight ADME genes with significant changes in the LPH environment to describe the prognostic features of pan-cancer. Lower risk scores calculated by the model were associated with better prognosis in 25 types of tumors, with a p-value of less than 0.05. The LPH environment was found to reduce the overall expression value of model genes, which could decrease the death risk of tumor prognosis. Additionally, it is found that the low-risk group had a higher degree of T cell infiltration based on immune infiltration analysis. Finally, drug exploration led to the identification of three potential model-regulating drugs. Overall, the study provided a new approach to construct a pan-cancer survival prognosis model based on ADME genes from the perspective of LPH and offered new ideas for future tumor prognosis research.

## Introduction

Improving the survival of patients with pan-cancer is a critical issue that tumor treatment faces at present^1,2^, and the main therapeutic tools currently used in clinical practice include immunotherapy^3,4^, adjuvant chemotherapy^5,6^, and cell base inactivation techniques^7^. However, these approaches still have some limitations, including decreasing the survival life span of healthy cells^8^, causing cellular damage^9^, and inducing severe adverse effects^8^, such as nausea and hair loss. Therefore, new therapeutic approaches that can improve tumor survival need to be further explored.

Recent studies have suggested that low press and hypoxia (LPH) environment may have beneficial effects on the prognosis of some tumors^10^. Several epidemiological researches have also proposed that long-term exposure to hypoxic environments may reduce tumor mortality^11,12^, such as the proportion of tumor deaths in Tibet was found to be the lowest among the 31 provinces in China mainland^13^. However, current works on the mechanisms of LPH to improve survival and inhibit tumorigenesis still have some limitations, as they are mostly only based on theoretical research such as iron transport and glucose transport hypotheses^14-16^, while in-depth studies on the effect of LPH environment on tumor survival at the level of population data are lacking. Meanwhile, some researches have indicated that LPH has effective regulatory impacts on some molecular markers^17,18^, for example, it has been shown that HIF factors in the hypoxia-sensing pathway, as crucial transcriptional regulators^19,20^, are involved in the regulation of erythrocyte metabolism^20^, which increases the transport of iron ions^14,16^. It is well known that iron metabolism is closely related to tumorigenesis, growth, and metastasis^21,22^, which reminds us of the key role that hypoxia and iron metabolism may play in cancer research.

Drug absorption, distribution, metabolism, and excretion (ADME) is a class of substances involved in the metabolism, transport, and detoxification of various endogens as well as exogenous substances in the human body^23^. The above as substrates of ADME may influence the process of cancer development and progression through direct effects, such as DNA damage, control of growth signaling pathways, and other phenomena, thus having a greater impact on patient survival. Therefore, exploring the prognostic relationship between cancer and ADME genes that are both LPH-responsive and associated with iron metabolism could provide a theoretical basis for LPH-assisted improvement of tumor prognosis.

## Method

### Data collection and process

Our research was conducted based on The Cancer Genome Atlas (TCGA) and tumor alterations relevant for genomics-driven therapy (TARGET) on the USCS Xena Server (https://xena.ucsc.edu/)^24^. The data for the TCGA/TARGET project came from the UCSC RNA-seq Compendium, samples of which were analyzed using RSEM (RNA-Seq by Expectation-Maximization). This study covered tumor data from the TCGA and TARGET databases of USCS Xena Server for more than 33 types of cancer, and we only employed its tumor samples (N = 10,137) to conduct gene expression analysis, after excluding cell line samples, metastatic samples, and null samples. We also included their gender information, overall survival status (OS), and overall survival time (OS time). In order to avoid the interference of short OS time, we excluded patients lived shorter than 60 days, and included gene (at least 20% gene with an expression value of 0) and Sample (No clinical features, more than 50% expression value is 0). The final sample size of cancer patients for survival analysis was 8,628.

### Specific genes screening and identification

Based on the MsigDB database, a collection of 38 biological pathway genes related to iron metabolism processes ^25^, consisting of 616 genes (**Table S1**), was included in the studies of Homo sapiens. Hypoxia-associated gene collections were obtained using the “Hypoxia” keyword in the H-gene collection (HALLMARK), which included 200 associated genes from 87 gene collections (**Table S2**). Afterward, 16 marker genes overlapping in the two collections were obtained using Venn analysis (**Table S3**). To screen strongly correlated drug metabolizing molecules, the expression profiles of the 298 ADME genes mentioned above were subjected to Spearman network construction and WGCNA network construction with the expression profiles of the 16 candidates, respectively ^23,26-28^. ADME genes with Spearmen correlation coefficients > |0.4| and weights of WGCNA edges > |0.07| were included. Univariate COX regression was conducted to further select the genes significantly associated with OS.

The study utilized the GEO dataset GSE103927 (AltitudeOmics) to identify ADME genes that demonstrated significant changes in a hypoxic environment by conducting ANOVA. GSE103927 involved analyzing the gene expression of mountaineers who ascended from 0-meter altitude into 5260-meters altitude and performed one iteration of the process. This study provided insights into the gene expression profile characteristics of the human body in an extreme hypoxic environment, making it a valuable contribution to the field of research on the effects of hypoxia on gene expression^29-31^, and could be beneficial for us to explore the alteration of drug metabolism gene expression profile in such environment. The above gene screening process is demonstrated in the flow chart in Fig 1A. To validate the model, we selected three expression profiles from the GEO dataset of two types of cancer with responsive clinical prognostic information. These included pancreatic adenocarcinoma, for which the GSE28735 and GSE57495 datasets were utilized, as well as gastric cancer, for which the GSE84437 dataset was used.

**Fig 1.**
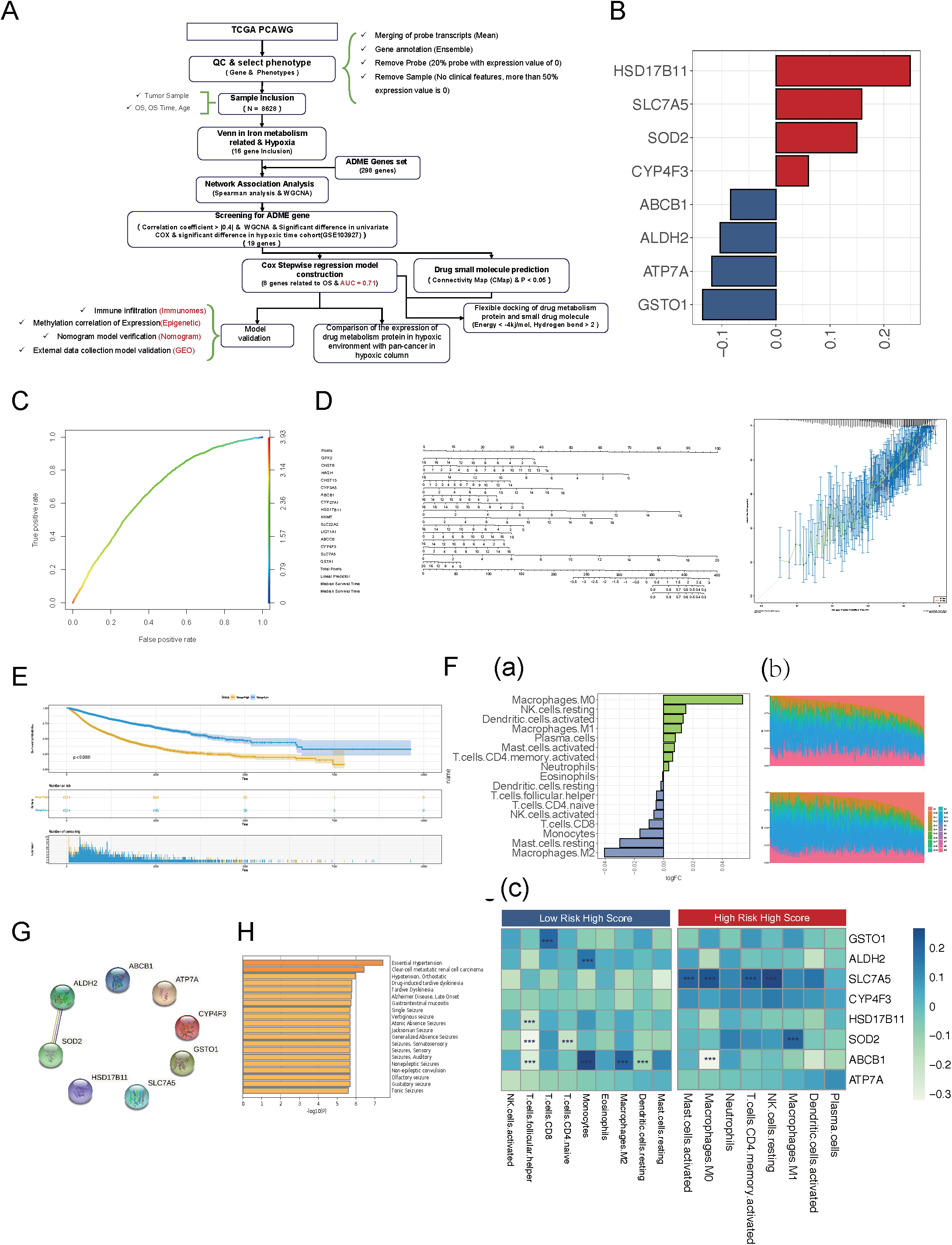
Selection of drug metabolism-related genes and prognostic model construction. A: Flow chart of data inclusion and statistical analysis; B: Risk prediction model coefficients for drug metabolism genes, where red represents positive and blue represents negative; C: 10-fold validation AUC; D: Nomogram graph that calibrated prediction scales for tumor OS using the 8 model genes at 3 years (C-index = 0.68) and 5 years (C-index = 0.68); E: Survival analysis prognostic curves of ADME in pan-cancer to distinguish the High- and the Low-Risk groups; F-(a): Difference in immune infiltration scores between the High-Risk and the Low-Risk groups, where the red bars represented high immune scores in the High-Risk group and the blue bars represented the ones in the Low-Risk group; F-(b): Distribution of immune infiltration scores in 22 immune cells, where the horizontal coordinates represented TCGA samples and the vertical coordinates represented scores; F-(c): Correlation between immune infiltration scores and expression of 8 ADME model genes; G: Protein-to-protein interaction network among the 8 model genes; H: Disease-related enrichment analysis for 8 model genes.

### ADME prognostic model construction

We used stepwise multivariate cox regression to further screen the candidate genes and finalize the tumor prognostic model with the following equation:

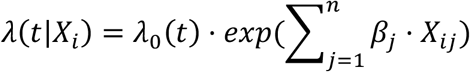

the predictive accuracy of the model was evaluated based on 10-fold AUC values and Nomogram^32^.

In the formula, *β*_*j*_ represented the fitting coefficient of the *j*th gene of the prognostic model, *X*_*i*_ denoted the expression value of the *j*th gene of *i*th sample, and *n* is the number of genes included in the prognostic model. Using this model, we classified the pan-cancer population into the High-Risk Group and the Low-Risk Group by model *Median* (*feature*_*sample*_)^8^. To compare the expression profile differences between the two groups, we used the Mann-Whitney U test for independent samples, and the degree of difference between the two groups was compared using log-transformed fold change (LogFC). One-way analysis of variance (ANOVA) was used for the comparison of more than two groups. We estimated survival curves using the Kaplan-Meier method and used the log-rank test for statistical comparisons.

To evaluate the level of response to the hypoxic environment, we explored the magnitude of hypoxic response scores in different risk groups by calculating the magnitude of ES scores in tumor samples in the hypoxic response pathway^33^ based on tumor expression profile data using the GSVA algorithm ^34^, results of which were evaluated by LogFC. Secondly, to explore whether the tumor immune microenvironment has differences in immune cell composition in different risk groups, the study used the CIBERSORT method ^35^ to estimate the abundance of immune infiltration in tumor patients using 22 immune cells. What’s more, considering that epigenetics can have a large impact on drug metabolism and resistance in cancer ^36^, this study performed the analysis of differences between risk groups using the Bumphunter algorithm ^37^ based on the epigenetic data of model genes (UCSC Xena Methylation 450K, https://pancanatlas.xenahubs.net). Afterward, cis-trans genes were also explored using the corresponding transcriptome profiles with methylation profiles ^38^.

All the processes were conducted with R (v 4.0.2). The significance level was set at 0.05 unless otherwise stated, and the multiple testing correction was adjusted by FDR (* denoted p-value ≤ 0.05; ** denoted p-value ≤ 0.01; *** denoted p-value ≤ 0.001; NS denoted p-value > 0.05).

### Small molecule prediction of drug metabolism

Based on the expression profile data of model ADME genes, we identified a set of small molecule drugs associated with the expression regulation of these genes ^39,40^, by using the L1000 platform of the Qure module (LINCS L1000 Chemical Perturbations 2021) ^41^ in CMAP’s small drug molecule prediction website (https://clue.io; https://maayanlab.cloud/). By searching for small molecules that reverse the expression profile of specific genes, drugs associated with the expression profile of the model ADME gene in the pan-cancer disease state were revealed (Z score < -4) ^39^, and small molecule drugs regulating each gene were identified using single-gene drug prediction (https://maayanlab.cloud/DGB/) ^42^. Subsequently, we construct the association networks between small molecule drugs and eight model genes using the STITCH v5.0 (http://stitch-beta.embl.de/) database ^43^, where the parameters of the networks were set as follows: The parameters were set as follows: organism = homo sapiens; active prediction methods = databases, Predictions and Experiments minimum required interaction score = medium confidence (0.400); max number of interactors to show = none (query proteins only); and view setting = evidence view. Drugs in the network that have interactions with model genes were selected for the follow-up analyses.

For the small molecule drugs with cancer therapeutic significance mentioned in the prediction section above, we performed flexible docking of drugs with model ADME genes, where the docking was performed using the algorithm provided by AutoDockTools 4.0 ^44^ (Binding energy < -4kj/mol, hydrogen bonds > 1). Afterward, the drug/cell line browser was used to explore the sensitivity of drug molecules in different cells, and the adverse drug reaction database was applied to investigate the negative reactions of drug molecules to ensure the stability of the drug administration process.

## Result

### Pan-ADME model

The Altitude Omics cohort, consisting of cancer-free individuals exposed to short- to long-term and repeated LPH at an altitude of 5800 meters, was analyzed to investigate the effect of LPH on the expression profiles of ADME genes. Of the 298 ADME genes identified to respond to LPH, we focused on the 19 genes that exhibited significant changes in expression under LPH and were associated with genes related to hypoxia and iron metabolism (**Table S1**). We selected eight of these genes to build a pan-cancer prognostic model (**Fig 1B)**, which demonstrated good predictive performance with an AUC of 0.71 (**Fig 1C**). A nomogram plot was utilized as a standard scale for tumor prognosis evaluation, the results supported the accuracy of the pan-ADME model in predicting the 3- and 5-year survival status of pan-cancer patients. (**Fig 1D**, C-index = 0.68). The pan-ADME model was established as follows:

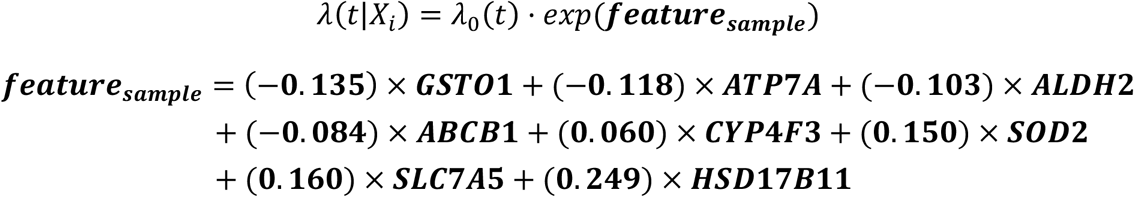

Our prognostic model exhibited strong performance in distinguishing individuals with high or low risk of death. We classified individuals with pan-cancer into High-Risk and Low-Risk groups using the ADME median value of the ADME model’s features, and evaluated the model effect on pan-cancer overall survival (OS). The results showed that the Low-Risk group had a significantly longer survival time compared with the High-Risk group with a p-value smaller than 0.001 (**Fig 1E**).

These selected ADME genes were further proved reasonable by immune infiltration analysis, on the consideration of the common sense that cancer is closely related to the immune status of the human body. Comparing the High-Risk and Low-Risk groups in the Pan-ADME model, we observed significant differences in 17 of 22 immune cell states (**Fig 1F-(a), Table S6**). The distribution of immune infiltration scores in 22 immune cells also showed differences in the High- and Low-Risk groups (**Fig 1F-(b)**). Notably, the T cell population was significantly higher in the Low-Risk group. Expression profiles of 5 ADME genes had significant correlations with the abundance of immune infiltration (**Fig 1F-(c)**), Table S6, p-value < 0.01, LogFC > |0.2|). The high abundance of follicular helper T-cells (Tfh) in the low-risk group was negatively correlated with 7 ADME genes, 3 of which were statistically significant (HSD17B11, SOD2, ABCB1; Table S6).

We conducted the Protein-Protein Interaction Networks (PPI) analysis and observed only ALDH2 has a protein interaction relationship with SOD2 (**Fig 1G**, Score= 0.589). This result indicated that most ADME genes did not interact with each other, which made it possible for us to improve the prognosis of cancer patients through single gene regulation. In order to explore the expression profiles and related characteristics of the 8 genes, the disease-related enrichment analysis was also performed, which supported their potential mechanism of essential hypertension (**Fig 1H**).

### ADME model in different types of cancer

Given that different types of cancer exhibit distinct characteristics in the tumor microenvironment, gene expression patterns may also differ across various cancer types. We utilized 8 candidate genes to construct prognostic models in 33 types of cancers, respectively. The results indicated that the prognosis status was significantly different among 25 (76% of 33) cancers (**Fig 2A**). To validate our findings, three expression profiles of two cancers (PAAD and STAD) with responsive clinical prognostic information from the GEO dataset were selected, on which applied our ADME genes to construct prognostic models. Their results also showed significant differences between the High-Risk and the Low-Risk groups, proving that AMDE gene expression can indeed have prognostic impacts in these cancers (**Fig 2B**).

**Fig 2.**
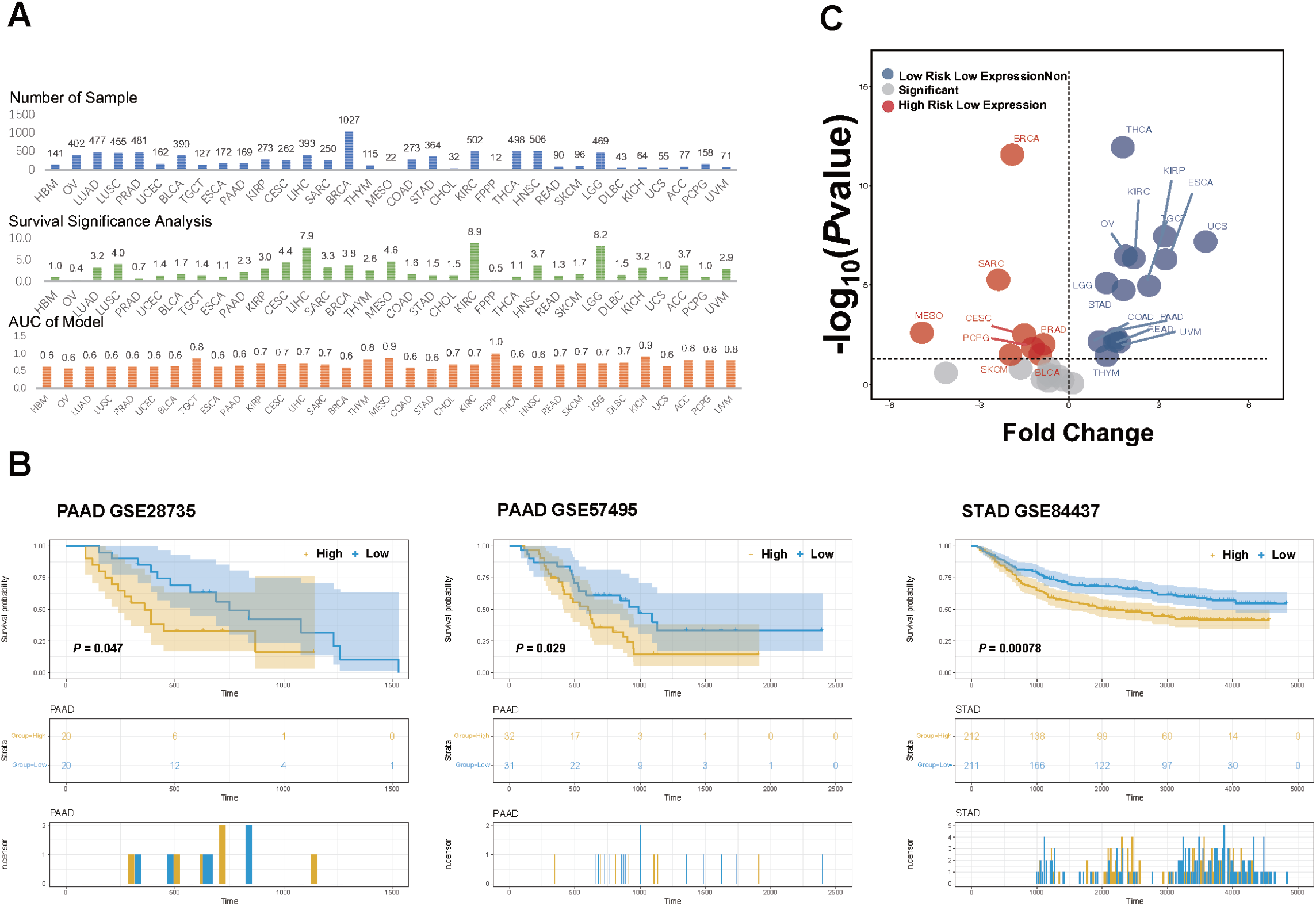
The total expression level of ADME genes in prognostic models in different cancers. A: Sample size, survival significance, and model AUC of 8 ADME gene models in 33 cancers (mean AUC=0.70); B: Volcano plot of the total expression values of ADME genes between the High- and Low-Risk groups in 33 cancers (logFC > 0 indicated that the total expression value of the Low-Risk group is lower and logFC < 0 indicated that the expression of the Low-Risk group is higher); C: Survival validation was performed in a collection of 3 expression profiles in 2 cancers (PAAD, STAD).

For all 33 kinds of cancers, we separately divided the samples into the High- and Low-Risk groups depending on their independent ADME models, and then compared the expression differences between groups. We observed the total expression values differed in 22 cancers, of which 14 cancers had significantly low expression levels in the Low-Risk group (logFC > 0, **Fig 2C**). Based on the prognostic model’s AUC results, we have tentatively demonstrated that the intervention of LPH could potentially decrease or improve the survival of certain tumors by reducing the total expression of ADME genes. And in subsequent analyses, we focused solely on cancers that met the following criteria: a) significant differences in OS between the risk groups; b) significant differences in total expression values between the risk groups; c) prediction AUC greater than 0.7; d) sample size larger than 100. Nine types of cancer fulfilled these criteria, including TGCT, KIRP, CESC, LIHC, SARC, THYM, MESO, KIRC, and LGG.

### Impact of LPH on cancer prognosis

We utilized the Altitude Omics cohort to investigate the response of 8 ADME genes under LPH (Altitude Omics cohort, **Fig 3A**, our findings revealed that the total expression profile of these model genes exhibited a strong LPH response capability (ANOVA p-value < 0.05) in the expression profiling study of the Altitude Omics cohort. Specifically, we observed a significant decrease in the total expression of the model genes following long-term (Time > day 16) and repeated (Re-day 7 and Re-day 21) LPH (**Fig 3B**). Then, we presented the box plots depicting the expression levels of 8 ADME genes at different time points, which revealed three distinct patterns of expression changes during LPH, including: (a) genes with reduced expression in long-term LPH: SOD2, ALDH2, HSD17B11, GSTO1, and ATP7A; (b) genes with reduced expression in short-term LPH: CYP4F3 and ABCB1; and (c) genes with increased expression in short-term LPH: SLC7A5 (**Fig 3C**).

**Fig 3.**
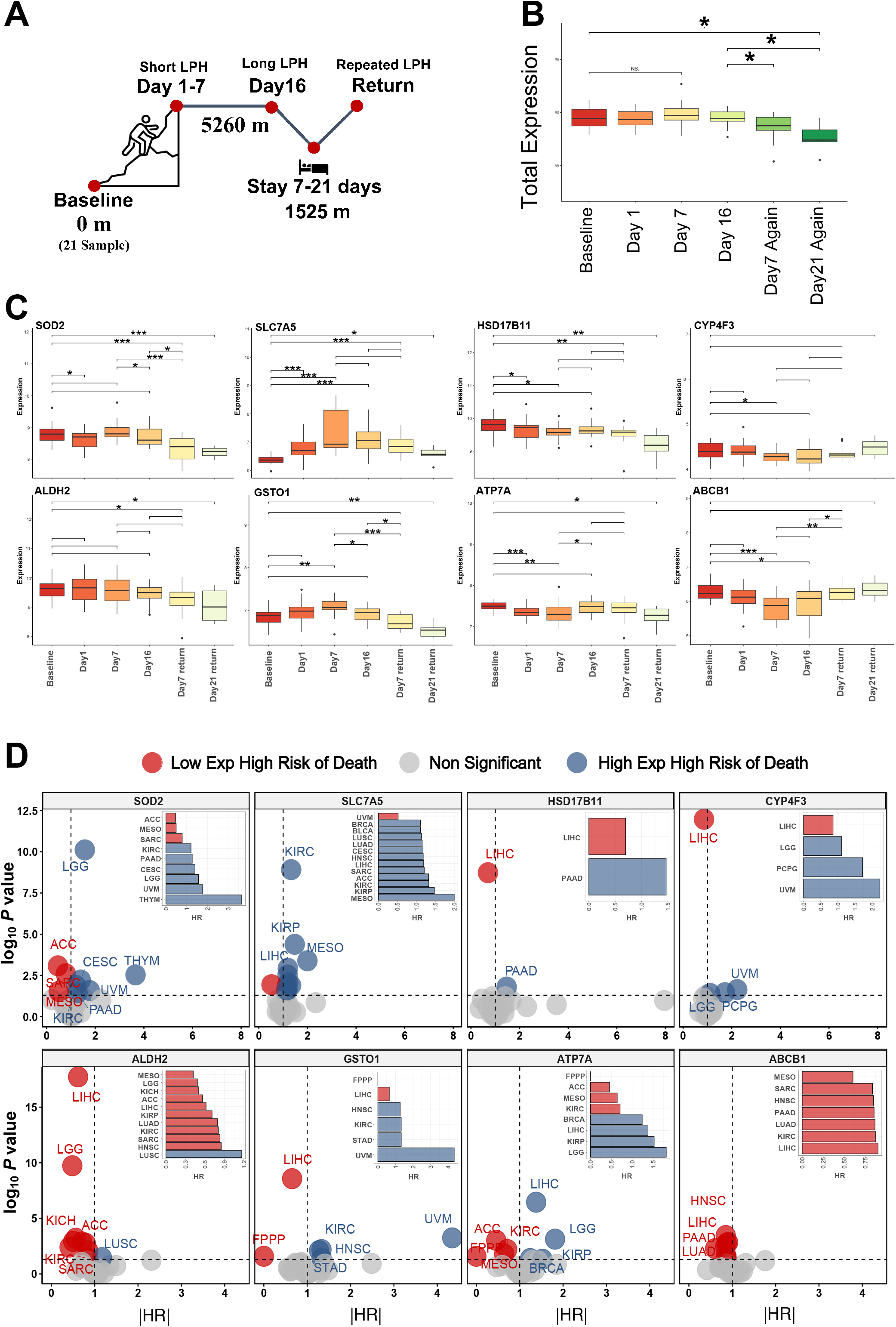
ADME prognostic model of single gene in different cancers. A: Description of AltitudeOmics cohort study; B: Trend of the total expression profile of 8 ADME model genes in the procedure of LPH; C: Trends of the expression profile of 8 genes during LPH separately; D: OS regressed on the single gene in 33 types of cancers, where HR>1 (blue bars) indicated the Low-Risk group with low expression values.

We have previously identified 9 tumors that are sensitive to a hypoxic environment, the OS of which is affected by the level of gene expression. In this part, we were interested in the effect of single-gene expression of the 8 ADME genes on the survival of each kind of tumor in the context of LPH, which might suggest the potential of each single gene as an independent molecular target.

We constructed single-cancer single-gene prognostic models and divided cancer patients into High- and Low-Risk groups and then compared the differences in expression values between the two groups (**Fig 3D**). Our focus was on the 7 genes exhibiting the first two patterns of expression changes, namely reduced expression in long-term or short-term LPH, and identified cancers with lower expression levels in the Low-Risk group (denoted as blue bars in **Fig 3D**). For example, SOD2 and GSTO1 exhibited low expression favoring OS in four (LGG, CESE, TYHM, KIRC) and one (KIRC) cancers, separately. In addition, ATP7A was helpful for patient survival in four cancers (LIHC, KIRP, LGG) as well as the low expression of CYP4F3 was beneficial to OS in LGG. In conclusion, we found that significant low expression of some genes under LPH condition may increase the survival of patients with tumors (CYP4F3, ATP7A, SOD2 for LGG; SOD2, GSTO1 for KIRC; ATP7A for LIHC; ATP7A for KIPR; SOD2 for CESE; SOD2 for TYHM). These findings suggested that downregulating different combinations of genes could potentially improve the survival of corresponding tumors.

### Drug Target Discovery and Intervention Strategies

We have demonstrated that the response of the 8 ADME genes to LPH could potentially impact tumor prognosis. However, we observed that the response of these genes to LPH was not uniform, which may attenuate the overall interventional effect of LPH. Additionally, we noted that SLC7A5 exhibited a distinct trend of up-regulation in short-term LPH, which may have a detrimental effect on survival when other genes’ expression is reduced in hypoxic environments. To further investigate this, we explored potential causes for the observed decrease in SLC7A5 expression over a longer period and considered methylation as a potential mechanism. We identified 37 CpGs of 8 ADME genes that showed significantly different between the High- and Low-Risk groups in pan-cancer (**Table S4**), and 4 Hypermethylation CpGs of SLC7A5 exhibited cis-action (coefficient < 0). Notably, a 1stExon island regional locus cg07067659 showed a significant hypermethylation trend after repeated LPH exposure in the Altitude Omics cohort, which might reduce the expression of SLC7A5 (**Fig 4A-(a)**). However, this level of decline might not be sufficient, so we introduced the concept of transcriptional profile pharmacological modulation of model genes as a means of regulation.

**Fig 4.**
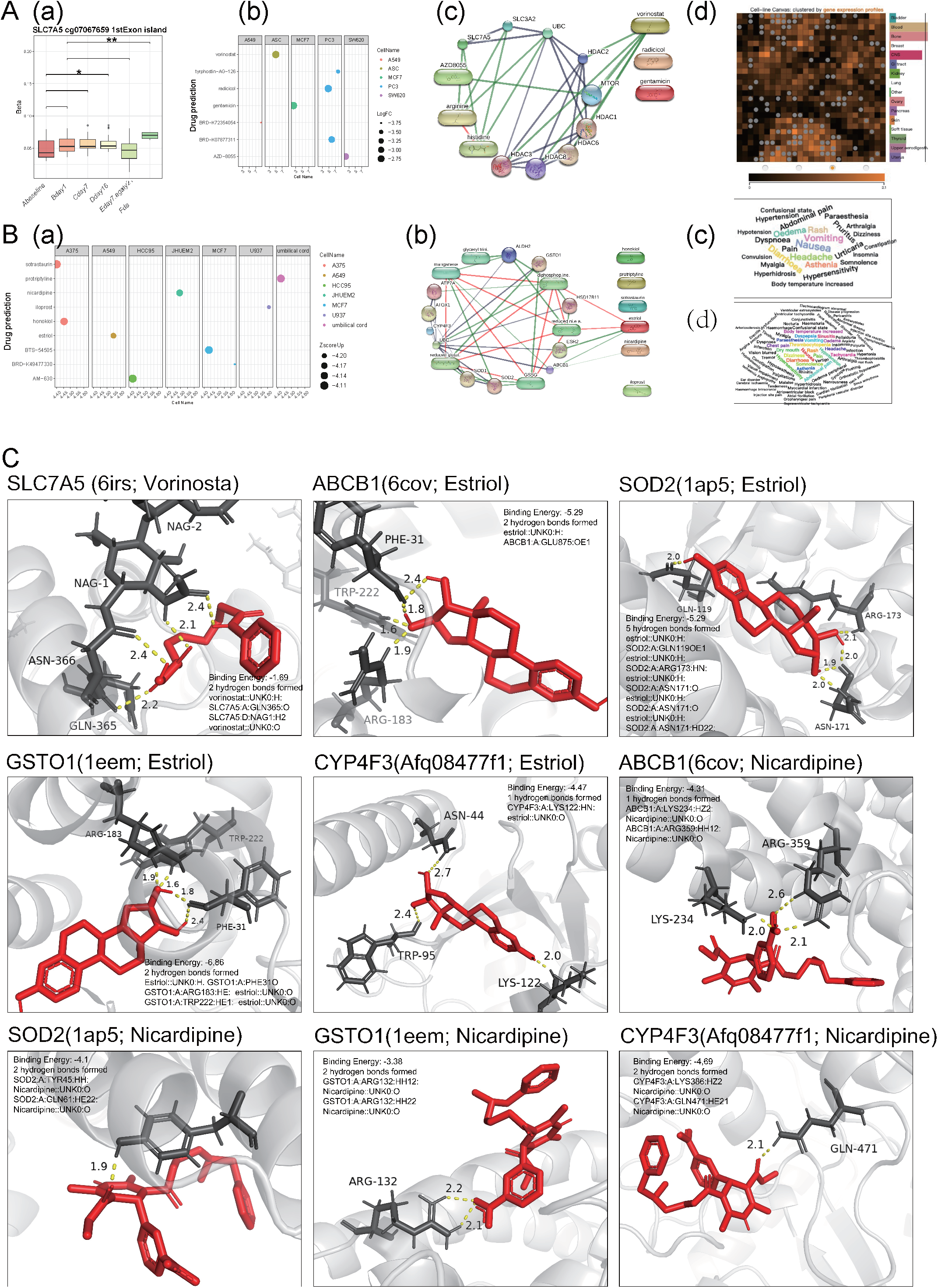
Drug molecule prediction with drug docking. A-(a): Change trend of methylation site cg07067659 in the AltitudeOmics cohort; A-(b): Small drug molecule enrichment using SLC7A5 (p-value < 0.05); A-(c): GDIN established for highly scored small drug molecules with SLC7A5; A-(d): Sensitivity test of Vorinosta in cells; B-(a): Small drug molecule enrichment using 7 other genes (Z-Score < -4, p-value < 0.05); B-(b): GDIN established for highly scored small drug molecules with 7 genes; B-(c): word cloud of major adverse effects of Nicardipine drug, where the larger the font and the closer to the center of the image the higher the probability of the adverse effects occurring; B-(d): word cloud of major adverse effects of Estriol drug; C: Drug docking (Energy <-4kj/mol and bonds >1) for each of the 3 drugs (Nicardipine, Estriol, Vorinosta) with corresponding ADME proteins.

After conducting a single-gene small molecule drug prediction for SLC7A5, we identified 7 drugs with significant differences (Fig 4A-(b), Table S8). Among them, Vorinosta showed promising results based on Gene and Drug Interaction Network (GDIN) analysis (Fig 4A-(c)) and drug docking analysis indicated a high binding energy with SLC7A5. However, there were also some hydrogen-built connections between them, which remained to be further explored (bonds = 2, **Fig 4C**). For Vorinosta, we found that it demonstrated significant sensitivity in various blood cells and bone cells (**Fig 4A-(d)**).

We conducted small molecule drug prediction analysis on the other 7 ADME genes, which suggested 27 potential small molecule drugs (Z-score < - 4, p-value < 0.01, **Fig 4B-(a), Table S7**). GDIN analysis suggested that 2 of the 27 drugs, estriol and nicardipine, might have interactions with 7 ADME expressed proteins (score > 0.400, **Fig 4B-(b)**). Based on drug docking, we further found that 4 ADME genes (GSTO1, ABCB1, SOD2, and CYP4F3) could be successfully docked with small drug molecules (**Fig 4C**). However, the common adverse reactions of the two drugs were mainly manifested as nausea and diarrhea (Fig 4B-(c)(d), Table S9).

## Discussion

In this study, we developed a prognostic model for pan-cancer utilizing ADME genes and investigated the potential of reducing cancer prognostic risk from a novel perspective via the adjuvant intervention of LPH. The results identified 8 genes as crucial targets for pan-cancer prediction with an AUC of 0.71, and 9 types of cancer might have reduced mortality risk under the influence of LPH. Further investigations unveiled that expression of ADME genes under LPH modified the immune infiltration status, thus impacting the tumor microenvironment. Moreover, we performed molecular drug prediction and docking of model genes, leading to the identification of three new targets for antitumor drugs. Overall, our study provided empirical evidence for the adjuvant effect of LPH on enhancing the survival of tumor patients and puts forth fresh perspectives for drug therapy.

We identified 8 ADME genes associated with LPH that may be beneficial in enhancing cancer survival. Among them, 8 genes except SLC7A5 were in a state of expression down-regulation at different time points of LPH intervention, while a significant low expression state was also observed in some types of cancer at low risk, and thus we claimed that the expression state of the 7 genes in these cancers is isotropic with low risk. ABCB1 and SOD2 were reported to be directly regulated by HIF-1, so they may play a key role in the adjuvant intervention of LPH. ABCB1 is regulated by HIF-1α in ovarian cancer cells ^45^, and lactate depletion in tumors can also significantly inhibit ABCB1 expression ^46^, while some studies have also suggested that its inhibitors can attenuate multidrug resistance in tumors after treatment ^47-50^. In addition, the increased expression of SOD2, an important antioxidant enzyme superoxide dismutase 2, stabilizes HIF-2α in an H2O2-dependent manner, promotes the expression of core stem cell genes (i.e., Oct4 and SOX2), and increases the cancer stem cell (CSC) subpopulation, tumorigenicity, and invasiveness of breast cancer cells ^51^. Conversely, the downregulation of SOD2 expression in cancer tissues attenuates cancer aggressiveness ^52-54^ and can help to sensitize cancer tissues to some chemotherapeutic agents ^55^. In addition, CYP4F3 has been found to be a possible prognostic influencer in lung cancer ^56^. ATP7A, on the other hand, is associated with neoangiogenesis as well as alleviating endothelial dysfunction ^57-59^, whose high expression levels may increase the survival risk of patients with advanced esophageal squamous cell carcinoma, ovarian cancer, etc. ^60-62^ The HSD17B11 gene plays an important role in androgen metabolism and its increased expression is associated with poorer survival in pancreatic cancer ^63^, as well as with advanced prostate cancer ^64^. GSTO1 expresses glutathione S-transferase Omega 1, whose expression is upregulated in several cancer cells and is associated with cancer progression ^65^. More lethally, it may promote the proliferation, migration, and invasion of non-small cell lung and breast cancer cells through the JAK/STAT3 signaling pathway and inhibit their apoptosis ^66,67^. ALDH2 is a widely recognized target for cancer therapy ^68,69^, and related studies have revealed that changes in its expression levels are associated with tumorigenesis and progression and differ between tumor types ^70^. Finally, for SLC7A5, it could express LAT1, which is seen as a molecular target for cancer diagnosis and therapy since the knockdown of LAT1 could suppress the proliferation of cancer cells and the growth of xenograft tumors^71^. Overall, the 8 model genes were shown to have regulatory roles in a variety of tumors and to be significantly associated with hypoxia-sensing pathways, as well as to be effective targets for tumor prognostic interventions in the LPH environment as ADME genes.

Furthermore, we explored the mechanisms of how the above ADME genes affect tumor prognosis. By immune infiltration analysis, we found that Tfh cells showed a negative correlation with the 7 ADME gene expression profiles. The Tfh cells secrete interleukin, IL-4, and IL-21, and thus promote B cell differentiation and antibody production during immunization^72-75^, which were found to be associated with increased and prolonged READ survival^76^. Our study indicated that ADME genes can be perturbed by the LPH environment and there was a higher degree of T-cell infiltration in the Low-Risk group, so we inferred that it could be possible that LPH influences T-cell infiltration through the response of the ADME genes to mediate the increase in OS of tumor patients.

In addition, small molecule drugs that could act with ADME genes have been suggested to be critical to complement LPH-assisted interventions ^77^. We identified 3 known small molecule drugs, Estriol, Nicardipine, and Vorinosta, that may be able to interact with 8 ADME genes. Some studies suggested the feasibility of these 3 drugs acting on ADME genes; for example, the downregulation of ABCB1^78^, SOD2^79^, ALDH2^79^, and GSTO1^80^ expression was found be associated with cisplatin resistance. Therefore, based on the expression profile of 8 ADME genes under LPH, we formulated a new therapeutic regimen such as KIRC in the LPH setting that could be combined with Estriol and Nicardipine drugs to have a significant impact on the prognosis of the tumor (**Table S10**). Although our study provides evidence of drug docking, the human in vivo environment is complex and protein interactions are complicated, so the specific drug effects need to be verified by relevant experiments at a later stage.

To the best of our knowledge, our study represents the first attempt to construct a pan-cancer prognostic prediction model using ADME-associated genes under LPH, to validate the model’s accuracy utilizing an external validation cohort, and to provide new insights into pharmacological interventions. Nonetheless, our research is constrained by the limited sample size of the study population; While we validated our model in various datasets, there is currently insufficient evidence on the regulation of ADME genes in hypoxic environments. Hence, expanding the corresponding population validation or conducting experimental validation could prove crucial. In future studies, we intend to explore the adjuvant effect of the LPH environment in greater depth, revealing its intervention effect on the survival of tumor patients.

## Author Contributions

**SX.Z**., **ZX.H. conceived and supervised the project. SX.Z**., **XL.W. conducted the data collection and processing. SX.Z**., **ZX.H**., **XX.H**., **XR.C**., **MZ.S. conducted the bioinformatics analysis, SX.Z**., **ZX.H**., **XX.H**., **XR.C**., **MZ.S. analyzed the resultsand wrote the manuscript, All the authors reviewed, commented and approved the manuscript**.

## Acknowledgments

We would like to appreciate the TCGA, GEO database for the availability of the data.

## References

1. Cheng, X., et al. Systematic Pan-Cancer Analysis Identifies TREM2 as an Immunological and Prognostic Biomarker. Front Immunol 12, 646523 (2021).

2. Chen, G., et al. GPC2 Is a Potential Diagnostic, Immunological, and Prognostic Biomarker in Pan-Cancer. Front Immunol 13, 857308 (2022).

3. Zhang, Y. & Zhang, Z. The history and advances in cancer immunotherapy: understanding the characteristics of tumor-infiltrating immune cells and their therapeutic implications. Cell Mol Immunol 17, 807–821 (2020).

4. Del Paggio, J.C. Immunotherapy: Cancer immunotherapy and the value of cure. Nat Rev Clin Oncol 15, 268–270 (2018).

5. Pondé, N.F., Zardavas, D. & Piccart, M. Progress in adjuvant systemic therapy for breast cancer. Nat Rev Clin Oncol 16, 27–44 (2019).

6. Morgensztern, D., et al. Adjuvant Chemotherapy for Patients with T2N0M0 NSCLC. J Thorac Oncol 11, 1729–1735 (2016).

7. Connor, M.J., et al. Cytoreductive treatment strategies for de novo metastatic prostate cancer. Nat Rev Clin Oncol 17, 168–182 (2020).

8. Muth, C.C. Chemotherapy and Hair Loss. Jama 317, 656 (2017).

9. Mokriani, S., Tukmechi, A., Harzandi, N. & Jabalameli, L. In vivo murine breast cancer targeting by magnetic iron nanoparticles involving L. GG cytoplasmic fraction. Iran J Basic Med Sci 24, 682–689 (2021).

10. Pérez-Herrero, E. & Fernández-Medarde, A. Advanced targeted therapies in cancer: Drug nanocarriers, the future of chemotherapy. Eur J Pharm Biopharm 93, 52–79 (2015).

11. Thiersch, M., Swenson, E.R., Haider, T. & Gassmann, M. Reduced cancer mortality at high altitude: The role of glucose, lipids, iron and physical activity. Exp Cell Res 356, 209–216 (2017).

12. Thiersch, M. & Swenson, E.R. High Altitude and Cancer Mortality. High Alt Med Biol 19, 116–123 (2018).

13. Lan, L., Zhao, F., Cai, Y., Wu, R.X. & Meng, Q. [Epidemiological analysis on mortality of cancer in China, 2015]. Zhonghua Liu Xing Bing Xue Za Zhi 39, 32–34 (2018).

14. Kautz, L., et al. Identification of erythroferrone as an erythroid regulator of iron metabolism. Nat Genet 46, 678–684 (2014).

15. Cook, J.D., Boy, E., Flowers, C. & Daroca Mdel, C. The influence of high-altitude living on body iron. Blood 106, 1441–1446 (2005).

16. Haase, V.H. Regulation of erythropoiesis by hypoxia-inducible factors. Blood Rev 27, 41–53 (2013).

17. Beall, C.M. Andean, Tibetan, and Ethiopian patterns of adaptation to high-altitude hypoxia. Integr Comp Biol 46, 18–24 (2006).

18. Cohen, E.B., Geck, R.C. & Toker, A. Metabolic pathway alterations in microvascular endothelial cells in response to hypoxia. PLoS One 15, e0232072 (2020).

19. Ke, Q. & Costa, M. Hypoxia-inducible factor-1 (HIF-1). Mol Pharmacol 70, 1469–1480 (2006).

20. Befani, C. & Liakos, P. The role of hypoxia-inducible factor-2 alpha in angiogenesis. J Cell Physiol 233, 9087–9098 (2018).

21. Wang, Y., Yu, L., Ding, J. & Chen, Y. Iron Metabolism in Cancer. Int J Mol Sci 20(2018).

22. Li, Y., et al. Hypobaric hypoxia regulates iron metabolism in rats. J Cell Biochem 120, 14076–14087 (2019).

23. Hu, D.G., Mackenzie, P.I., Nair, P.C., McKinnon, R.A. & Meech, R. The Expression Profiles of ADME Genes in Human Cancers and Their Associations with Clinical Outcomes. Cancers (Basel) 12(2020).

24. Goldman, M.J., et al. Visualizing and interpreting cancer genomics data via the Xena platform. Nat Biotechnol 38, 675–678 (2020).

25. Silgado-Guzmán, D.F., et al. Characterization of ADME Gene Variation in Colombian Population by Exome Sequencing. Front Pharmacol 13, 931531 (2022).

26. Hu, D.G., Marri, S., McKinnon, R.A., Mackenzie, P.I. & Meech, R. Deregulation of the Genes that Are Involved in Drug Absorption, Distribution, Metabolism, and Excretion in Hepatocellular Carcinoma. J Pharmacol Exp Ther 368, 363–381 (2019).

27. Jittikoon, J., et al. Comparison of genetic variation in drug ADME-related genes in Thais with Caucasian, African and Asian HapMap populations. J Hum Genet 61, 119–127 (2016).

28. Klein, K., et al. A New Panel-Based Next-Generation Sequencing Method for ADME Genes Reveals Novel Associations of Common and Rare Variants With Expression in a Human Liver Cohort. Front Genet 10, 7 (2019).

29. Bourdillon, N., et al. AltitudeOmics: Baroreflex Sensitivity During Acclimatization to 5,260 m. Front Physiol 9, 767 (2018).

30. Fan, J.L., et al. AltitudeOmics: Resetting of Cerebrovascular CO2 Reactivity Following Acclimatization to High Altitude. Front Physiol 6, 394 (2015).

31. Ryan, B.J., et al. AltitudeOmics: rapid hemoglobin mass alterations with early acclimatization to and de-acclimatization from 5260 m in healthy humans. PLoS One 9, e108788 (2014).

32. Iasonos, A., Schrag, D., Raj, G.V. & Panageas, K.S. How to build and interpret a nomogram for cancer prognosis. J Clin Oncol 26, 1364–1370 (2008).

33. Eustace, A., et al. A 26-gene hypoxia signature predicts benefit from hypoxia-modifying therapy in laryngeal cancer but not bladder cancer. Clin Cancer Res 19, 4879–4888 (2013).

34. Hänzelmann, S., Castelo, R. & Guinney, J. GSVA: gene set variation analysis for microarray and RNA-seq data. BMC Bioinformatics 14, 7 (2013).

35. Chen, B., Khodadoust, M.S., Liu, C.L., Newman, A.M. & Alizadeh, A.A. Profiling Tumor Infiltrating Immune Cells with CIBERSORT. Methods Mol Biol 1711, 243–259 (2018).

36. Thakur, C. & Chen, F. Connections between metabolism and epigenetics in cancers. Semin Cancer Biol 57, 52–58 (2019).

37. Morris, T.J., et al. ChAMP: 450k Chip Analysis Methylation Pipeline. Bioinformatics 30, 428–430 (2014).

38. Zhang, Y. & Chen, L. [DNA methylation and non-small cell lung cancer]. Zhongguo Fei Ai Za Zhi 13, 821–826 (2010).

39. Yang, P.M., Chou, C.J., Tseng, S.H. & Hung, C.F. Bioinformatics and in vitro experimental analyses identify the selective therapeutic potential of interferon gamma and apigenin against cervical squamous cell carcinoma and adenocarcinoma. Oncotarget 8, 46145–46162 (2017).

40. Chakarov, S., et al. Two distinct interstitial macrophage populations coexist across tissues in specific subtissular niches. Science 363(2019).

41. Subramanian, A., et al. A Next Generation Connectivity Map: L1000 Platform and the First 1,000,000 Profiles. Cell 171, 1437-1452.e1417 (2017).

42. Wang, Z., et al. Drug Gene Budger (DGB): an application for ranking drugs to modulate a specific gene based on transcriptomic signatures. Bioinformatics 35, 1247–1248 (2019).

43. Kuhn, M., et al. STITCH 4: integration of protein-chemical interactions with user data. Nucleic Acids Res 42, D401–407 (2014).

44. Forli, S., et al. Computational protein-ligand docking and virtual drug screening with the AutoDock suite. Nat Protoc 11, 905–919 (2016).

45. Parmakhtiar, B., Burger, R.A., Kim, J.H. & Fruehauf, J.P. HIF Inactivation of p53 in Ovarian Cancer Can Be Reversed by Topotecan, Restoring Cisplatin and Paclitaxel Sensitivity. Mol Cancer Res 17, 1675–1686 (2019).

46. Wang, J.W., et al. A Self-Driven Bioreactor Based on Bacterium-Metal-Organic Framework Biohybrids for Boosting Chemotherapy via Cyclic Lactate Catabolism. ACS Nano 15, 17870–17884 (2021).

47. Liu, C., Xing, W., Yu, H., Zhang, W. & Si, T. ABCB1 and ABCG2 restricts the efficacy of gedatolisib (PF-05212384), a PI3K inhibitor in colorectal cancer cells. Cancer Cell Int 21, 108 (2021).

48. Zhang, Y., et al. Poziotinib Inhibits the Efflux Activity of the ABCB1 and ABCG2 Transporters and the Expression of the ABCG2 Transporter Protein in Multidrug Resistant Colon Cancer Cells. Cancers (Basel) 12(2020).

49. Besse, A., et al. Carfilzomib resistance due to ABCB1/MDR1 overexpression is overcome by nelfinavir and lopinavir in multiple myeloma. Leukemia 32, 391–401 (2018).

50. Goto, S., Kawabata, T. & Li, T.S. Enhanced Expression of ABCB1 and Nrf2 in CD133-Positive Cancer Stem Cells Associates with Doxorubicin Resistance. Stem Cells Int 2020, 8868849 (2020).

51. Bonini, M., et al. 122 - Acetylation activates an alternative function of SOD2 as a stemness factor in breast cancer. Free Radical Biology and Medicine 128, S62–S63 (2018).

52. Piskounova, E., et al. Oxidative stress inhibits distant metastasis by human melanoma cells. Nature 527, 186–191 (2015).

53. Buchheit, C.L., Weigel, K.J. & Schafer, Z.T. Cancer cell survival during detachment from the ECM: multiple barriers to tumour progression. Nat Rev Cancer 14, 632–641 (2014).

54. Yen, H.C., et al. Alterations of the levels of primary antioxidant enzymes in different grades of human astrocytoma tissues. Free Radic Res 52, 856–871 (2018).

55. Zhou, J. & Du, Y. Acquisition of resistance of pancreatic cancer cells to 2-methoxyestradiol is associated with the upregulation of manganese superoxide dismutase. Mol Cancer Res 10, 768–777 (2012).

56. Zhao, M., et al. Identification of immune-related gene signature predicting survival in the tumor microenvironment of lung adenocarcinoma. Immunogenetics 72, 455–465 (2020).

57. Ash, D., et al. The P-type ATPase transporter ATP7A promotes angiogenesis by limiting autophagic degradation of VEGFR2. Nat Commun 12, 3091 (2021).

58. Sudhahar, V., et al. Caveolin-1 stabilizes ATP7A, a copper transporter for extracellular SOD, in vascular tissue to maintain endothelial function. Am J Physiol Cell Physiol 319, C933–c944 (2020).

59. Ashino, T., et al. Copper transporter ATP7A interacts with IQGAP1, a Rac1 binding scaffolding protein: role in PDGF-induced VSMC migration and vascular remodeling. Am J Physiol Cell Physiol 315, C850–c862 (2018).

60. Li, Z.H., Lu, X., Li, S.W., Chen, J.T. & Jia, J. Expression of ATP7A in esophageal squamous cell carcinoma (ESCC) and its clinical significance. Int J Clin Exp Pathol 12, 3521–3525 (2019).

61. Lukanović, D., Herzog, M., Kobal, B. & Černe, K. The contribution of copper efflux transporters ATP7A and ATP7B to chemoresistance and personalized medicine in ovarian cancer. Biomed Pharmacother 129, 110401 (2020).

62. Yu, Z., Cao, W., Ren, Y., Zhang, Q. & Liu, J. ATPase copper transporter A, negatively regulated by miR-148a-3p, contributes to cisplatin resistance in breast cancer cells. Clin Transl Med 10, 57–73 (2020).

63. Bai, R., Rebelo, A., Kleeff, J. & Sunami, Y. Identification of prognostic lipid droplet-associated genes in pancreatic cancer patients via bioinformatics analysis. Lipids Health Dis 20, 58 (2021).

64. Nakamura, Y., Suzuki, T., Arai, Y. & Sasano, H. 17beta-hydroxysteroid dehydrogenase type 11 (Pan1b) expression in human prostate cancer. Neoplasma 56, 317–320 (2009).

65. Djukic, T., et al. Upregulated glutathione transferase omega-1 correlates with progression of urinary bladder carcinoma. Redox Rep 22, 486–492 (2017).

66. Wang, K., Zhang, F.L. & Jia, W. Glutathione S-transferase ω 1 promotes the proliferation, migration and invasion, and inhibits the apoptosis of non-small cell lung cancer cells, via the JAK/STAT3 signaling pathway. Mol Med Rep 23(2021).

67. Manupati, K., et al. Glutathione S-transferase omega 1 inhibition activates JNK-mediated apoptotic response in breast cancer stem cells. Febs j 286, 2167–2192 (2019).

68. Shao, C., et al. Essential role of aldehyde dehydrogenase 1A3 for the maintenance of non-small cell lung cancer stem cells is associated with the STAT3 pathway. Clin Cancer Res 20, 4154–4166 (2014).

69. Liu, X., et al. Targeting ALDH1A1 by disulfiram/copper complex inhibits non-small cell lung cancer recurrence driven by ALDH-positive cancer stem cells. Oncotarget 7, 58516–58530 (2016).

70. Zhang, H. & Fu, L. The role of ALDH2 in tumorigenesis and tumor progression: Targeting ALDH2 as a potential cancer treatment. Acta Pharm Sin B 11, 1400–1411 (2021).

71. Elorza, A., et al. HIF2α acts as an mTORC1 activator through the amino acid carrier SLC7A5. Mol Cell 48, 681–691 (2012).

72. Zhou, D.M., et al. The role of follicular T helper cells in patients with malignant lymphoid disease. Hematology 22, 412–418 (2017).

73. Victora, G.D. & Nussenzweig, M.C. Germinal Centers. Annu Rev Immunol 40, 413–442 (2022).

74. Tellier, J. & Nutt, S.L. The unique features of follicular T cell subsets. Cell Mol Life Sci 70, 4771–4784 (2013).

75. Crotty, S. T Follicular Helper Cell Biology: A Decade of Discovery and Diseases. Immunity 50, 1132–1148 (2019).

76. Bindea, G., et al. Spatiotemporal dynamics of intratumoral immune cells reveal the immune landscape in human cancer. Immunity 39, 782–795 (2013).

77. Yu, A.M., et al. Regulation of drug metabolism and toxicity by multiple factors of genetics, epigenetics, lncRNAs, gut microbiota, and diseases: a meeting report of the 21(st) International Symposium on Microsomes and Drug Oxidations (MDO). Acta Pharm Sin B 7, 241–248 (2017).

78. Huang, Y.W., et al. Amphiregulin promotes cisplatin chemoresistance by upregulating ABCB1 expression in human chondrosarcoma. Aging (Albany NY) 12, 9475–9488 (2020).

79. Kim, J., Chen, C.H., Yang, J. & Mochly-Rosen, D. Aldehyde dehydrogenase 2*2 knock-in mice show increased reactive oxygen species production in response to cisplatin treatment. J Biomed Sci 24, 33 (2017).

80. Tsuboi, K., et al. Potent and selective inhibitors of glutathione S-transferase omega 1 that impair cancer drug resistance. J Am Chem Soc 133, 16605–16616 (2011).

